# *Hox* gene expression in postmetamorphic juveniles of the brachiopod *Terebratalia transversa*

**DOI:** 10.1101/449488

**Authors:** Ludwik Gąsiorowski, Andreas Hejnol

## Abstract

**Background:** *Hox* genes encode a family of homeodomain containing transcription factors that are clustered together on chromosomes of many Bilateria. Some bilaterian lineages express these genes during embryogenesis in spatial and/or temporal order according to their arrangement in the cluster, a phenomenon referred to as collinearity. Expression of *Hox* genes is well studied during embryonic and larval development of numerous species; however, relatively few studies focus on the comparison of pre- and postmetamorphic expression of *Hox* genes in animals with biphasic life cycle. Recently, the expression of *Hox* genes was described for embryos and larvae of *Terebratalia transversa*, a rhynchonelliformean brachiopod, which possesses distinct metamorphosis from planktonic larvae to sessile juvenile. During premetamorphic development, *T. transversa* does not exhibit spatial collinearity and several of its *Hox* genes are recruited for the morphogenesis of novel structures. In our study we determined expression of Hox genes in post-metamorphic juveniles of *T. transversa* in order to examine metamorphosis-related changes of expression patterns and to test if Hox genes are expressed in the spatially collinear way in the postmetamorphic juveniles.

**Results:** *Hox* genes are expressed in a spatially non-collinear manner in juveniles, generally showing similar patterns as ones observed in competent larvae: genes *labial* and *Post1* are expressed in chaetae-related structures, *sex comb reduced* in the shell forming epithelium, whereas *Lox5* and *Lox4* in dorso-posterior epidermis. After metamorphosis expression of genes *proboscipedia, Hox3, Deformed* and *Antennapedia* becomes restricted to, respectively, shell musculature, prospective hinge rudiments, and pedicle musculature and epidermis.

**Conclusions:** All developmental stages of *T. transversa*, including postmetamorphic juveniles, exhibit a spatial non-collinear *Hox* genes expression with only minor changes observed between pre- and postmetamorphic stages. Our results are concordant with morphological observation that metamorphosis in rhynchonelliformean brachiopods, despite being rapid, is rather gradual. The most drastic changes in *Hox* genes expression patterns observed during metamorphosis could be explained by the inversion of the mantle lobe, which relocates some of the more posterior larval structures into the anterior edge of the juveniles. Cooption of *Hox* genes for the morphogenesis of novel structures is even more pronounced in post-metamorphic brachiopods when compared to larvae.

## Background

*Hox* genes encode a family of conserved homeodomain transcription factors from the ANTP class, which by binding to regulatory DNA sequences can activate or suppress transcription of downstream genes (e.g. [1, 2]). *Hox* genes are present in genomes of almost all investigated animals (with exception of Porifera, Ctenophora and Placozoa [3–7]) and are hypothesized to represent a synapomorphy of the clade consisting of Cnidaria and Bilateria [4, 8–10]. In most of Bilaterians *Hox* genes are expressed during embryogenesis, being involved in antero-posterior (A-P) patterning of either the whole embryo or at least some of its developing organ systems (e.g. [1, 2, 11]). Interestingly, in the genomes of some animals, the *Hox* genes are clustered along the chromosomes in the same order as they are expressed along A-P axis, a phenomenon referred to as spatial collinearity [2, 11–13]. The clustering of *Hox* genes in the genome is hypothesized as a plesiomorphic feature of Bilateria (e.g. [13]), which however, went through extensive remodeling in some evolutionary lineages (e.g. [12, 14–22]. Yet, spatial collinearity can be preserved despite a disorganization or split of the ancestral *Hox* cluster (e.g. [14]), the situation for which the term trans-collinearity was coined by Duboule [12].

Initially the role of *Hox* genes has been studied in the developing embryo of *Drosophila melanogaster* [23], later supplemented by the data from other insects, vertebrates and nematodes [24–26]. Recent advance of molecular and bioinformatic techniques allowed the investigation of *Hox* gene expression in the embryos and larvae of several non-model species, including e.g. xenacoelomorphs [16, 27, 28], hemichordates [29], onychophorans [30], tardigrades [31], rotifers [32], annelids [33–35], mollusks [36–40], nemerteans [41] and brachiopods [19], essentially increasing knowledge on diversity of Hox genes based patterning systems in Bilateria.

Many animals are characterized by an indirect life cycle in which embryos develop through a larval stage and subsequent metamorphosis, during which the larval body is reshaped into the adult one (e.g. [42, 43]). As larvae and adults can significantly differ in their morphology, the transition process might be quite dramatic and hence attracted attention of many researchers as one of the pivotal moments of the animal development [44–46]. Although the process of metamorphosis has puzzled numerous developmental biologists, there are relatively few studies regarding shifts of *Hox* gene expression accompanying it [15, 47–51]. In some animals both larvae and adults show canonical spatial collinearity, which often correlates with gradual type of metamorphosis. This can be exemplified by investigated annelid species, in which both life stages exhibit spatial collinearity of most of the *Hox* genes, yet there are shifts in the combinations of genes defining particular body regions before and after metamorphosis [47, 48]. On the other hand, in the other animals (especially those with the more pronounced metamorphosis) only one of the developmental stages exhibits canonical spatial collinearity of *Hox* genes expression, whereas the remaining stage shows either a non-collinear expression or does not express *Hox* genes at all. For instance, in the tunicate *Ciona intestinalis Hox* genes exhibit spatially collinear expression in the nervous system of larvae, whereas in juveniles only three posterior genes are expressed in the intestine [15]. Conversely in pilidiophoran nemertean *Micrura alaskensis* and in indirectly developing enteropneust *Schiozcardium californicum* the specialized larvae develop without expressing any of the Hox genes, which, in turn, are expressed in the canonical collinear way only in the rudiments of juvenile worms developing either inside larval body (pilidiophorans) or as the posterior extension of late larva (enteropneusts) [49, 50]. A somehow similar situation is found in the indirectly developing sea urchin *Strongylocentrotus purpuratus*, in which only two *Hox* genes (*Hox7 and Hox 11/13b*) take part in the larva formation, whereas the rudiments of adult animal, developing inside the larval body, show collinear expression of five *Hox* genes (*Hox7, Hox8, Hox9/10, Hox11/13a and Hox11/13b*) in the extra-axial mesoderm [51–54]. It is therefore evident, that metamorphosis-related shifts in *Hox* gene expression and function vary a lot from one animal clade to another, as a result of diverse evolutionary and developmental processes which shape the ontogeny of each particular group [55].

One of the animal groups with a distinct metamorphosis event are rhynchonelliformean brachiopods, represented by *Terebratalia transversa* for which Schiemann et al. recently described Hox genes expression in embryos and larvae [19]. Brachiopods, along with phoronids and possibly ectoprocts, constitute the clade Lophophorata (Fig.1 A, [56, 57]), which, together with e.g. annelids, mollusks, flatworms, nemerteans and rotifers, belongs to a large clade of protostome animals called Spiralia (Fig. 1A, [57–60]). Extant brachiopods are traditionally divided into three groups: Rhynchonelliformea, Craniiformea and Linguliiformea, the two latter forming sister clades [18, 57, 60, 61], historically united into group Inarticulata. As all brachiopods, adults of *T. transversa* are filter feeding animals with external anatomy superficially similar to bivalves – most of the body, including lophophore, a filtering organ, is enclosed in the two-valved shell, which covers the dorsal and ventral surfaces of the body. The clade Rhynchonelliformea is further characterized by the set of morphological features, including: posterior soft-tissued pedicle (by which animal attaches to the substrate), blind gut devoid of anus and articulated valve-hinge [62]. Additionally, rhynchonelliformean larvae differ from those found in other brachiopods by possessing three distinct body regions (Fig. 1B1) – anterior lobe, mantle lobe (bearing four chaetal sacs) and the most posterior pedicle lobe [62–66]. The rhynchonelliformean larva settles by adhering to the substrate with the posterior tip of the pedicle lobe (Fig. 1B2) and undergoes a specific metamorphosis, which in case of *T. transversa* is relatively rapid (few hours to one day [63]) and involves inversion of the mantle lobe (Fig. 1B2–3, [63, 67]). The latter results in profound relocation of some larval tissues – in competent larvae the mantle lobe partially covers the pedicle one and its chaetae projects posteriorly, after metamorphosis mantle lobe with chaetae projects anteriorly, its former interior surface becomes exposed and produce protegulum (the first rudiment of the shell), whereas its former exterior surface constitute walls of the mantle cavity of juvenile (Fig. 1B2–4) [63, 67, 68]. Therefore, the rapid transition from larvae to the juvenile involves profound reshaping of the entire body, which poses a question to which extent are those two stages are continuous [69].

**Fig. 1.**
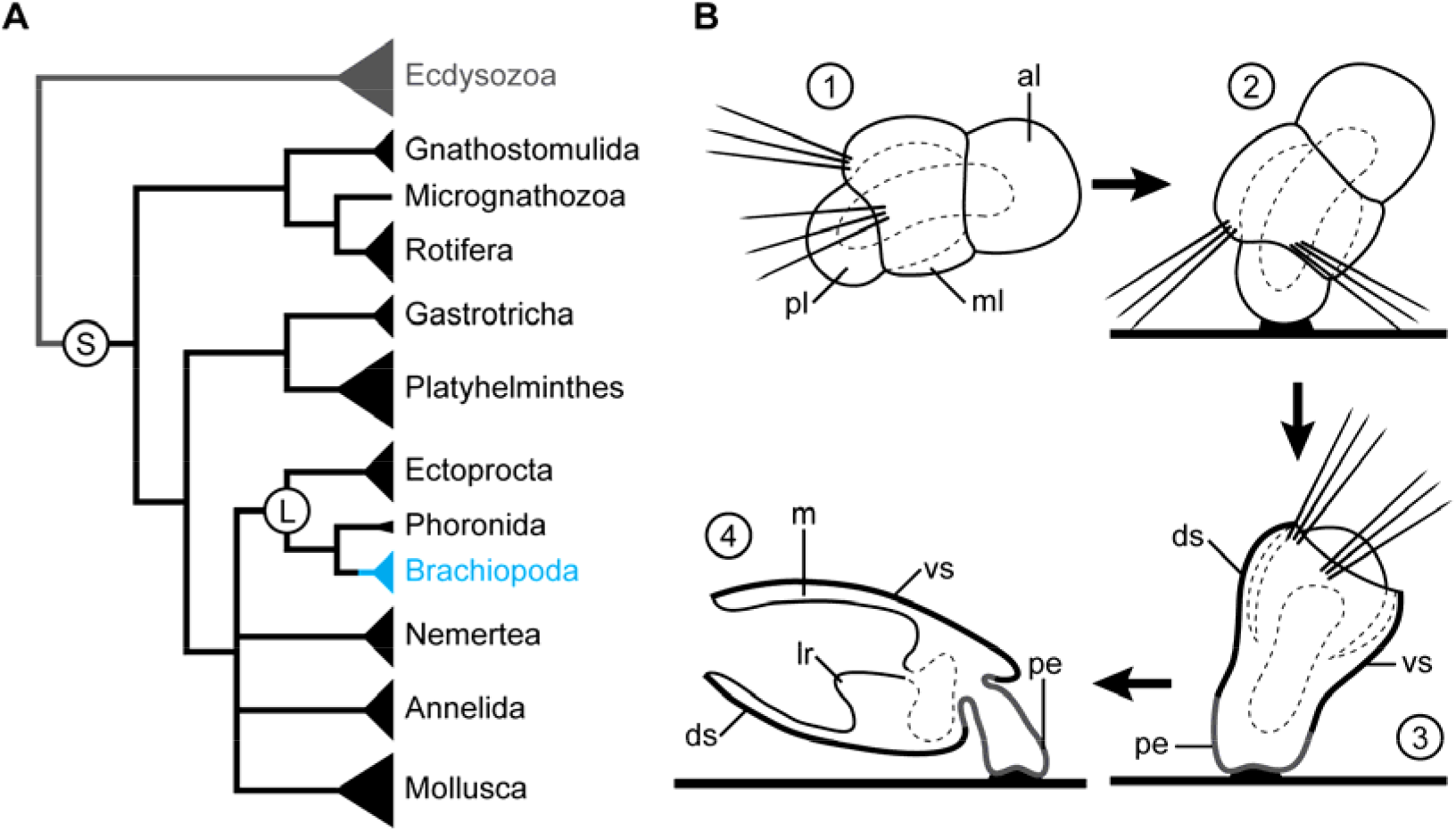
Phylogenetic position of Brachiopoda (A, based on Laumer et al 2015[57]) and metamorphosis of *Terebratalia transversa* (B, based on Freeman 1993 [67]). S stands for Spiralia, L for Lophophorata. 1. Competent planktonic larva, anterior to the right; 2. Larva settles on the substrate; 3. Inversion of the mantle lobe in the settled larva; 4 Juvenile; note that over the course of metamorphosis the internal surface of the larval mantle lobe becomes external, shell covered surface of juvenile animal, external surface of the mantle lobe becomes inner surface of the mantle, whereas anterior lobe contributes to the lophophore rudiment developing inside mantle cavity. Abbreviations: *al* larval anterior lobe, *ds* dorsal shell, *lr* lophophore rudiment, *m* mantle, *ml* larval mantle lobe, *pe* pedicle, *pl* larval pedicle lobe, *vs* ventral shell.

Schiemann et al. investigated genomic order of *Hox* genes of *T. transversa* and Hox genes expression in embryos and larvae of *T. transversa* and craniiformean Novocrania anomala [19]. *T. transversa* has a split *Hox* cluster comprising of 10 *Hox* genes in three independent parts. One scaffold contains two anterior *Hox* genes, labial (lab) and *proboscipedia* (*pb*). Second, longest part, contains genes *Hox3*, *deformed (Dfd), sex combs reduced* (Scr), *Lox5, antennapedia (Antp), Lox4* and *Post2*, whereas the most posterior gene *Post1* is located in the third independent scaffold [19]. A disorganization of the *Hox* cluster has also been reported for linguliformean brachiopod, *Lingula anatina*, in which although all *Hox* genes are in the single cluster *Post1, Post2, Lox4* and *Antp* has been translocated upstream to the *lab* [18]. In embryos and larvae of *T. transversa*, but also of craniiformean *N. anomala*, detected expression pattern of *Hox* genes does not show the canonical spatial collinearity [19]. However, as stated before, in some indirectly developing animals, larvae and juveniles can show collinear expression of *Hox* genes in patterning of one of life stages while other develops without evident Hox expression collinearity.

Therefore, in this study, we supplemented findings of Schiemann et al. (2017) by examination of the postmetamorphic *Hox* gene expression in *Terebratalia transversa* juveniles 2 days after metamorphosis. The main questions, which we were aiming to answer, were: 1) If and how is *Hox* gene expression pattern shifted during metamorphosis in rhynchonelliformean brachiopods? 2) Is there any staggered *Hox* gene expression along the A-P axis emerging after metamorphosis as a result of displacement and development of certain larval Anlagen?

## Results

### Description of *T. transversa* juvenile morphology

The existing knowledge on the detailed morphology of juvenile *T. transversa* is based mostly on confocal laser scanning microscopy (CLSM) investigation of musculature [69, 70], as well as transmission electron microscope (TEM) sections [63, 68, 71] and scanning electron microscopy (SEM) [68, 71, 72] of different developmental stages, including juveniles 1 day after metamorphosis [68, 69, 72], 4–5 days after metamorphosis [68, 70–72] and older than one week after metamorphosis [68, 69, 71, 72]. Therefore, to facilitate interpretation of our gene expression results [73], we examined morphology of the juveniles two days after metamorphosis utilizing light microscopy (LM) and CLSM combined with DAPI, phalloidin and immunohistochemical stainings (with primary antibodies against tyrosinated and acetylated tubulin).

Two days after metamorphosis juveniles of *T. transversa* already resemble the adult animal in their general shape (Fig. 2A–C). The body is clearly divided into main part covered by the two-valved juvenile shell and a posterior pedicle (*pe*, Fig. 2A–C), by which juvenile is attached to the substrate.

**Fig. 2.**
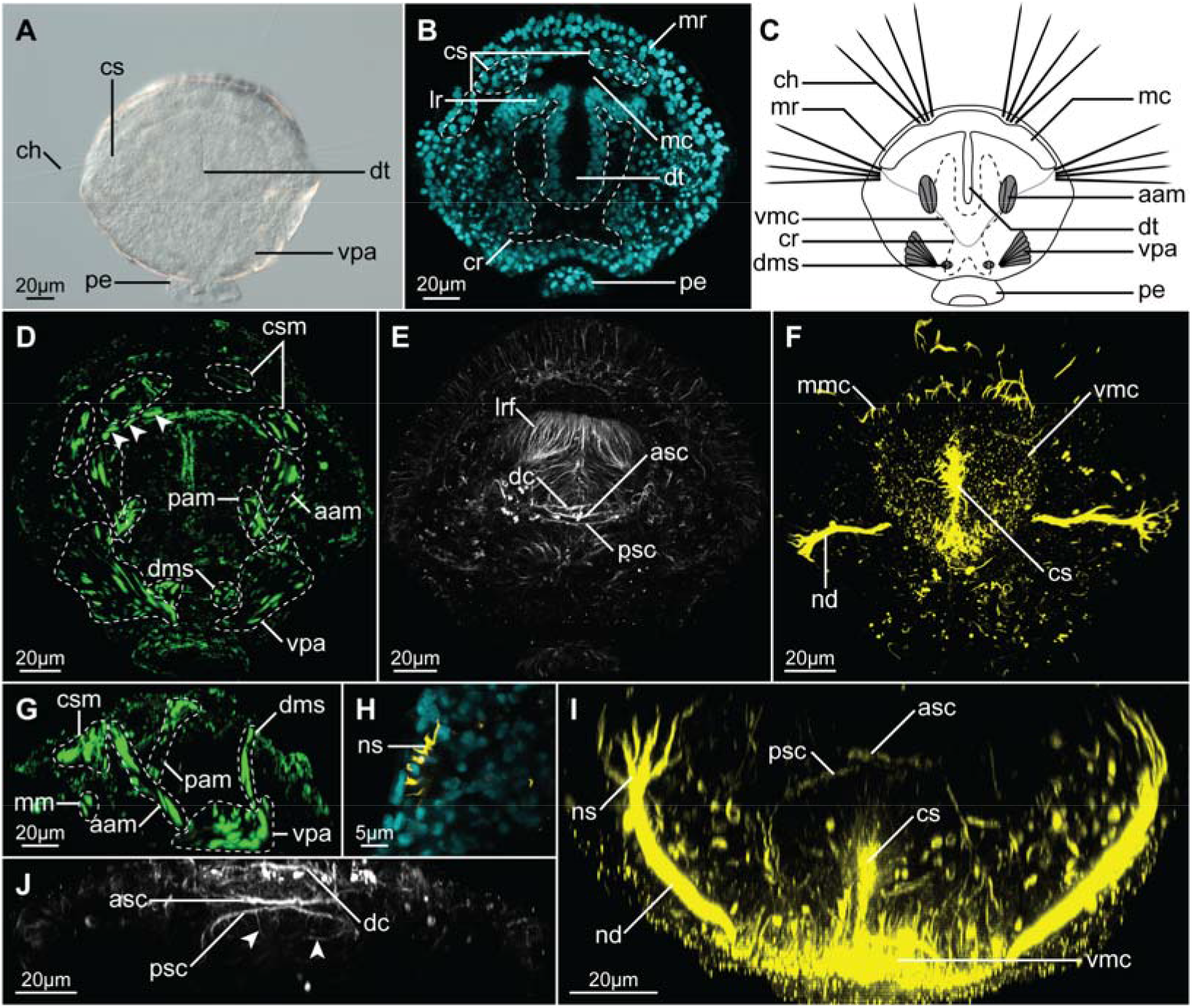
Morphology of the juvenile of *Terebratalia transversa* two days after metamorphosis, visualized with light microscopy (A) and CLSM (B, D-I). A Micrograph of the entire animal. B Frontal section through the median part of animal, cell nuclei visualized with DAPI staining. C Schematic drawing of the anatomy of juvenile. D, G Musculature visualized with F-actin phalloidin staining, arrowheads in D points to the ventral mantle margin muscles. E, J tyrosinated tubulin-like immunoreactivity. F, I acetylated-tubulin-like immunoreactivity. H Transverse section through the nephrostome, cell nuclei visualized with DAPI in cyan, acetylated tubulin-like immunoreactivity in yellow. A-F Dorso-ventral view, anterior to the top. G Lateral view, anterior to the left, dorsal to the top. I Virtual transverse section through the middle part of animal, dorsal to the top. J. Ventral, magnified view of the commissural region from E. Abbreviations: *aam* anterior shell adductor muscle, *asc* anterior supraoesophageal commissure, *ch* chaeta, *cr* coelom rudiment, *cs* chaetal sac, *csm* chaetal sac musculature, *dc* dorsal commissure, *dms* shell diductor muscle, *dt* digestive tract, *lr* lophophore rudiment, *lrf* fibers in the lophophore rudiment, *mc* mantle cavity, *mm* mantle margin musculature, *mmc* mantle margin ciliation, *mr* mantle margin, *nd* nephroduct, *ns* nephrostome, *pam* posterior shell adductor muscle, *pe* pedicle, *psc* posterior supraoesophageal commissure, *vmc* ventral mantle cavity, *vpa* ventral pedicle adjustor muscle.

Anteriorly, the shell is lined with mantle margin (*mr*, Fig. 2B, C), where tissues responsible for secretion of prospective adult shell are localized (Stricker and Reed 1985a). Phalloidin staining revealed presence of developing mantle margin muscles (arrowheads Fig. 2D; *mm*, Figs. 2G and 3C, C’), which have been already described for the older juveniles [69, 70]. Additionally, four chaetal sacs (*cs*, Fig. 2A, B), associated with degenerating larval musculature (csm, Figs. 2D, G; 3D, D’) [69], are embedded in the mantle margin, one pair dorso-medially and another in the more lateral position, which respectively protrudes numerous chaetae (*ch*, Fig. 2A, C) anteriorly and laterally.

**Fig. 3.**
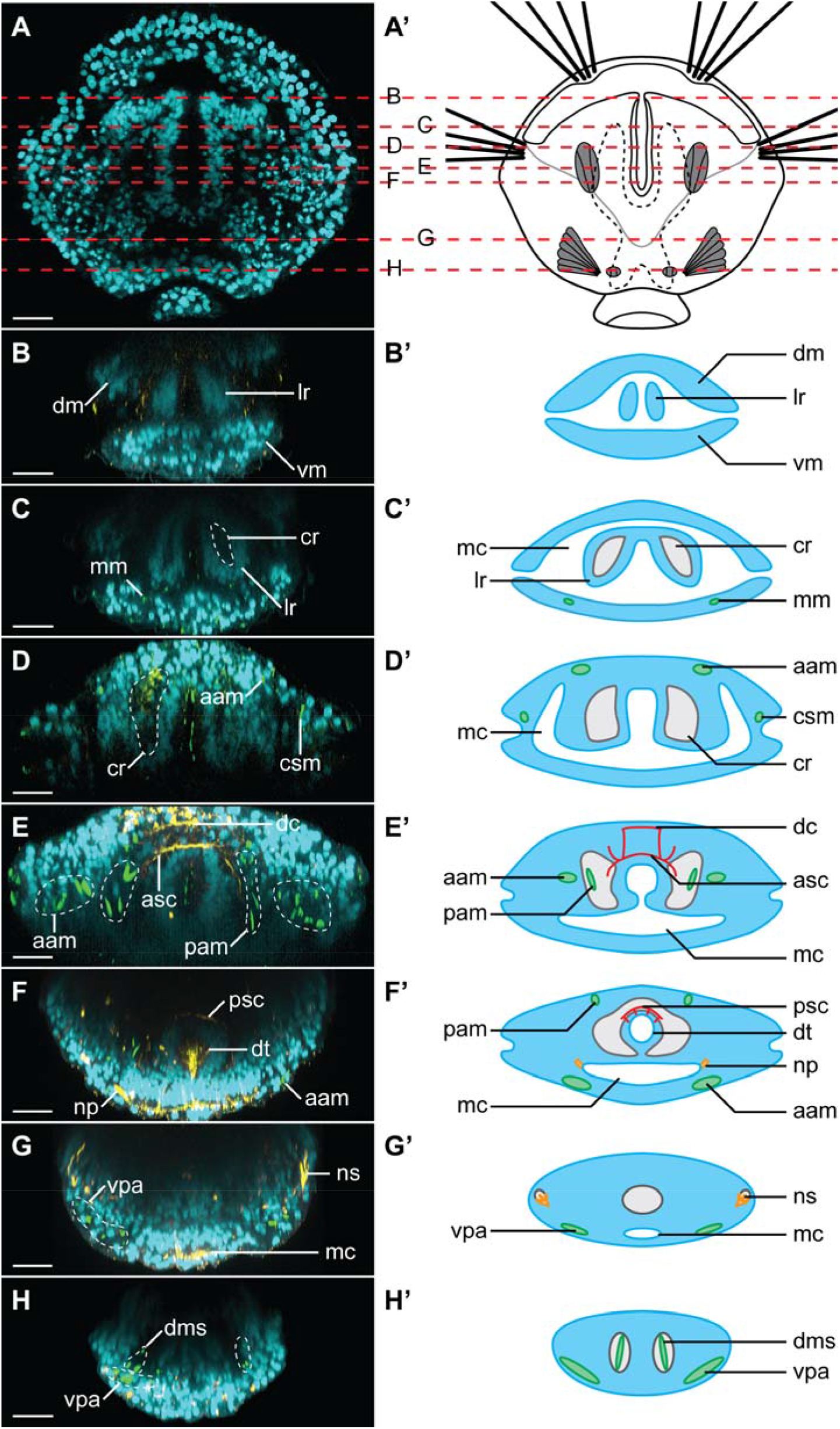
Transverse sections through *Terebratalia transversa* juvenile (two days after metamorphosis), showing detailed morphology of the animal. Each panel consists of CLSM image (left) and schematic representation (right). Cell nuclei visualized with DAPI in cyan, acetylated-tubulin like immunoreactivity (F and G) and tyrosinated tubulin-like immunoreactivity (E) in yellow, F-actin phalloidin staining in green. A Juvenile in dorso-ventral view, anterior to the top, dashed red lines indicates sections shown on the subsequent panels. B-H Virtual transverse sections through the juvenile, as indicated on A, dorsal to the top on all panels, on the right panels outline of the body is depicted in light blue, musculature in green, body cavities in grey, nervous system in red and excretory organs in orange. Abbreviations: *aam* anterior shell adductor muscle, *asc* anterior supraoesophageal commissure, *cr* coelom rudiment, *csm* chaetal sac muscle, *dc* dorsal commissure, *dm* dorsal mantle, *dms* shell diductor muscle, *dt* digestive tract, *lr* lophophore rudiment, *mc* mantle cavity, *mm* mantle margin muscle, *np* nephropore, *ns* nephrostome, *pam* posterior shell adductor muscle, *psc* posterior supraoesophageal commissure, *vm* ventral mantle, *vpa* ventral pedicle adjustor muscle.

Optical sections through the animal show the narrow mantle cavity (*mc*, Figs. 2B, C; 3C’–G’), which expands ventro-medially to about two-thirds the length of animal body and is lined with ciliated cells (*vmc*, Fig. 2C, F, I). The remnant of the larval anterior lobe, from which the prospective lophophore will develop [65, 74], is situated inside the mantle cavity (*lr*, Figs. 2B; 3B, B’, C, C’). Posteriorly the lobe is connected with the dorsal mantle and ventrally it faces the extension of the mantle cavity (Fig. 3D, D’, E, E’). Medially the lobe is divided by the ciliated slit (*cs*, Fig. 2F, I), which anteriorly communicates with the mantle cavity through the ventral infold, a stomodeum (Fig. 3 C–E, C’–E’) and posteriorly continues as the tubular rudiment of digestive tract (*dt*, Figs. 2A–C; 3F, F’).

At this stage the lophophore rudiment is poorly developed and consists of two, scarcely ciliated lobes without tentacles (*lr*, Figs. 2B; 3B, B’, C, C’). The lobes are penetrated by numerous, fine tyrosinated-tubulin-like immunoreactive (tTLIR) fibers (*lrf*, Fig. 2E), some of which communicate with the nervous system and probably represent the developing innervation of the prospective lophophore.

The nervous system is inconspicuous. Its most prominent structures are two tTLIR and acetylated-tubulin like immunoreactive (aTLIR) commissures (anterior and posterior supraoesophageal commissures; respectively: *asc* and *psc*, Figs. 2E, I, J; 3E, E’, F, F’), positioned dorsally to the ciliated slit, more or less one third from its most posterior extremity. Some tTLIR and aTLIR fine fibers extend laterally from those commissures to lophophore rudiments and mantle tissues (including mantle margin and chaetal sacs). Few tTLIR fine neurites extends from posterior supraoesophageal commissure to the intestinal tissue (*arrowheads*, Fig. 2J; Fig. 3F’). Additional dorsal tTLIR commissure (*dc*, Figs. 2E, J; 3E, E’) connects with the anterior supraoesophageal commissure. The similar arrangement of nervous system in early juveniles of *T. transversa* has been reported based on immunostaining against serotonin [28].

Two prominent tTLIR and aTLIR longitudinal structures are present in the ventral part of the animal (*nd*, Fig. 2F, I), extending along the v entral surface from the mid-posterior region to the ventral part of the mantle cavity. Dorsally those structures have numerous finger-like projections (*ns*, Figs. 2H, I, 3G, G’), which contact nuclei-free regions (as revealed by DAPI staining, Fig. 2H). We suggest, that those structures represent metanephridia (composed of nephrostome and nephridial duct) of juveniles, which connects the developing coelom with the mantle cavity. Their form and position is similar to what has been described for the metanephridia of relatively closely related Terebratulina retusa [75]. Although metanephridia in brachiopods are considered to be responsible only for release of gametes and not for excretion[75], they are present (albeit initially as non-functional rudiments) already in the early juveniles of *N. anomala* [76]. It is possible that aTLIR structures described by Santagata [69] as larval protonephridia in *T. transversa* actually represent rudiments of the metanephridial ducts or nephrostomes which acquire their final form during or soon after metamorphosis.

The DAPI staining revealed an empty cavity inside the body of the juvenile with two pairs of anterior and posterior branches (cr, Figs. 2B, C; 3), which most probably represents the developing coelom in which some of the forming muscles are freely positioned (Fig. 3E, E’, H, H’) [77]. Its two anterior branches extend along digestive tract and penetrate the lophophore rudiment (Fig. 2B; 3C–F, C’–F’). A similar arrangement of the coelom in the lophophore rudiment of postmetamorphic juveniles has been described for relatively closely related rhynchonelliformean *Calloria inconspicua* [78].

In addition to the already mentioned musculature related with the mantle margin, we identified rudiments of all the muscle groups (pedicle adjustors, shell diductors as well as anterior and posterior shell adductors; respectively: vpa, dms, aam, pam, Figs. 2D, G; 3D–H, D’–H’) described for the older juveniles of *T. transversa* [70] with the only exception of lophophore-related tentacle muscles (which correlates with lack of lophophore tentacles 2 days after metamorphosis). In different specimens the particular groups of muscles were developed to different degree corroborating the observation of the extensive and rapid remodeling of muscular tissue in the postmetamorphic juveniles [69].

### *In situ* hybridization of *Hox* genes

The expression of the *Hox* genes in juvenile *Terebratalia transversa* (2 days after metamorphosis) was examined with colorimetric (CISH; Fig. 4) and fluorescent (FISH, Fig. 5) in *situ* hybridization. Generally, *Hox* genes in the juveniles of *Terebratalia transversa* are not expressed in a collinear way (Figs. 4–6).

**Fig. 4.**
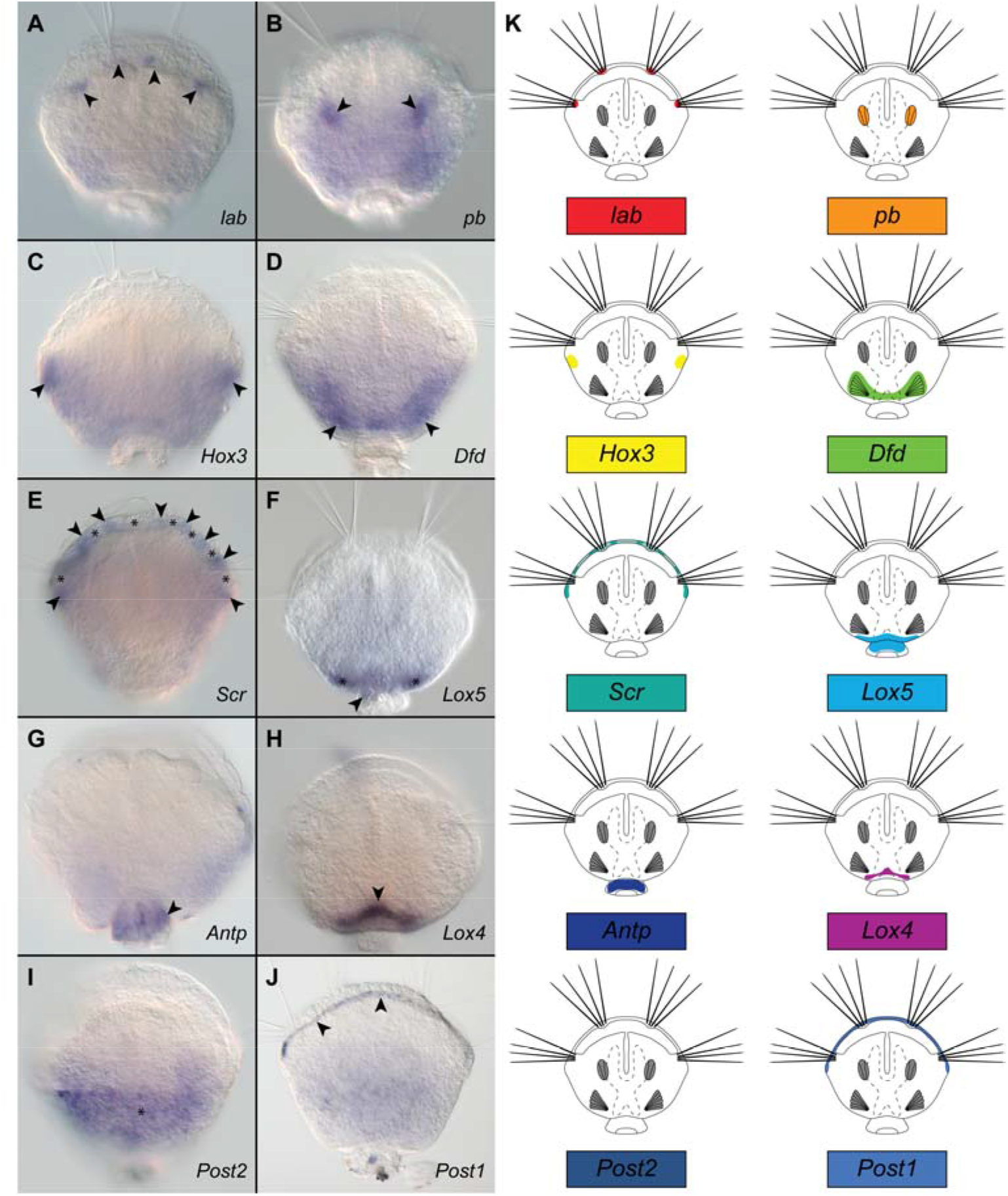
Whole-mount colorimetric *in situ* hybridization of the *Hox* genes (A–J) and schematic representation of expression patterns (K) in *Terebratalia transversa* postmetamorphic juveniles (two days after metamorphosis). All micrographs and drawings in dorso-ventral view, anterior to the top. For each plate (A–J) name of the hybridized gene is provided in the lower right corner. Particular structures in which each of the genes is expressed are indicated with arrowheads, asterisks and double arrowheads (see text for detailed explanation). Note that signal on I (asterisk) represents unspecific background (see text for details).

**Fig. 5.**
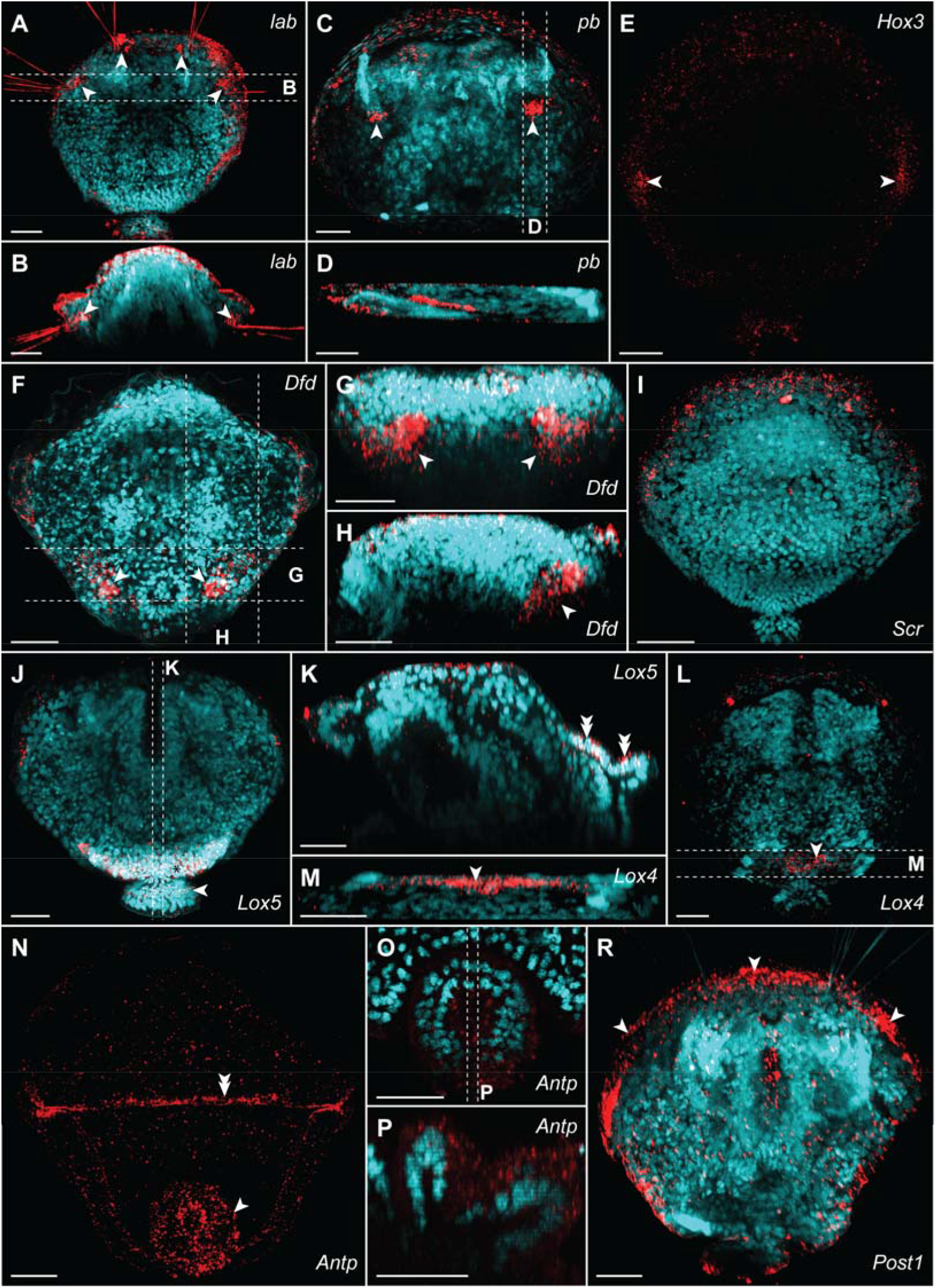
Whole-mount fluorescent *in situ* hybridization of the *Hox* genes (red) combined with DAPI staining of cell nuclei (cyan, with exception of E and N) in *Terebratalia transversa* postmetamorphic juveniles (two days after metamorphosis). For each plate name of the hybridized gene is provided in the right corner. Particular structures in which each of the genes is expressed are indicated with arrowheads, asterisks and double arrowheads (see text for detailed explanation). Expression of *lab* in chaetal sacs in dorso-ventral view (A) and virtual cross section (B); note autofluorescence of chaetae. Expression of *pb* in anterior shell adductor muscles in dorso-ventral view (C) and virtual parasagittal section (D). Expression of *Hox3* in dorso-ventral view (E). Expression of *Dfd* in ventral pedicle adjustor muscles in dorso-ventral view (F), virtual cross section (G) and parasagittal section (H). Expression of *Scr* in mantle margin (I). Expression of *Lox5* in dorso-posterior epidermis in dorso-ventral view (J) and virtual sagittal section (K). Expression of *Lox4* in dorso-posterior epidermis in dorso-ventral view (L) and virtual cross section (M). Expression of *Antp* in pedicle tissue in dorsoventral view (N), and virtual cross (O) and longitudinal (P) sections through pedicle. Expression of Post 2 in mantle margin, dorso-ventral view (R). Dashed lines with letters on A, C, F, J, L and O indicate section planes shown on respective plates. Scale bars on all images represent 20μm. Anterior to the top on A, C, E, F, I, J, L, N and R; dorsal to the top on B, D, G, H, K and M; anterior to the left on D, H and K.

**Fig. 6.**
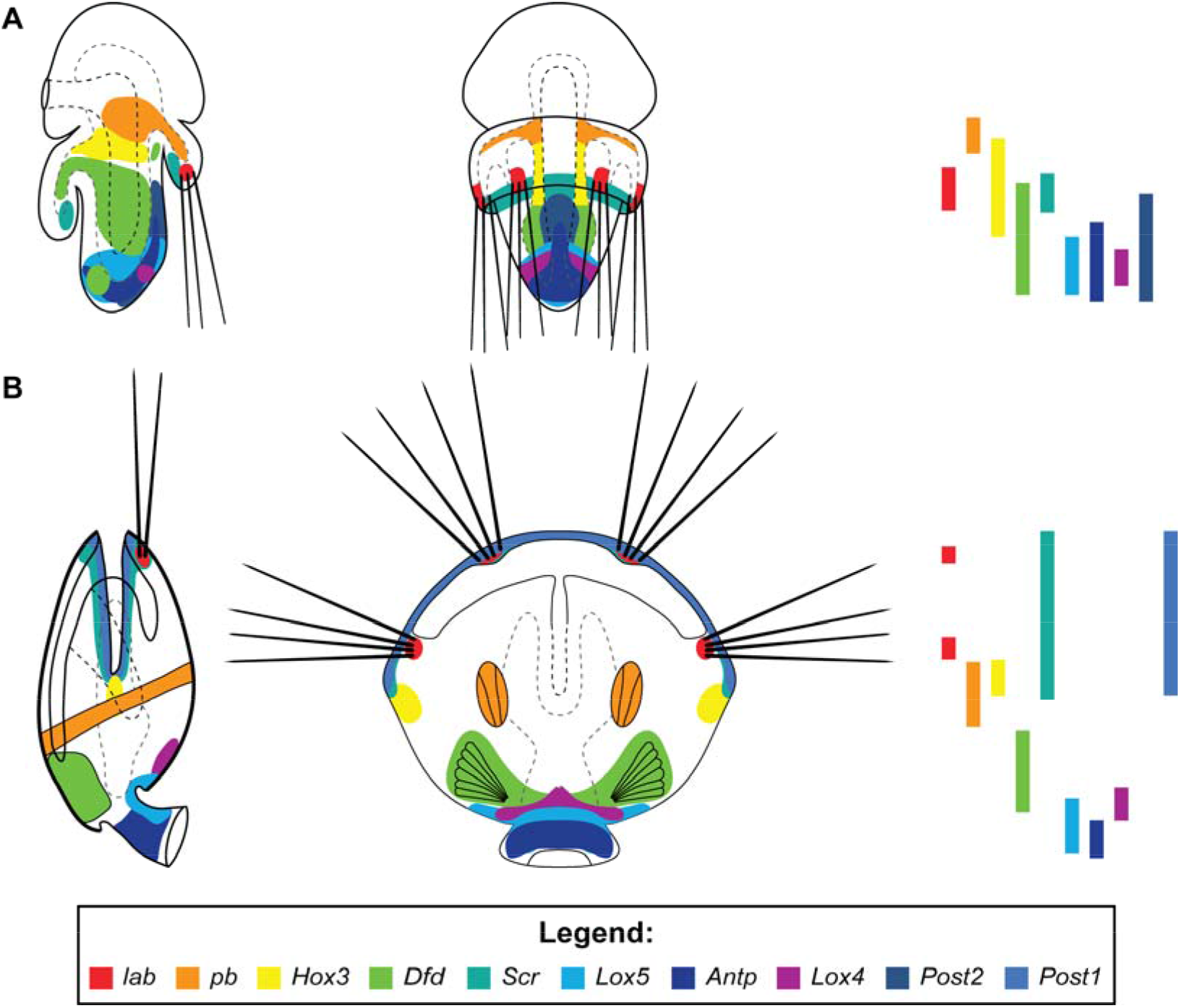
Comparison of the *Hox* genes expression between the late competent larva (A, based on Schiemann et al. 2017[19]) and juvenile (B) of *Terebratalia transversa*. Animals are shown in the dorso-ventral view (right panels) and in the lateral view with dorsal to the right (left panels). Anterior to the top on all panels. Bars on the right show antero-posterior *Hox* gene expression gradients in each developmental stage.

The most anterior *Hox* gene lab is expressed in two bilaterally paired domains at the mantle margin, which correspond to the larval chaetal sacs (arrowheads, Figs. 4A; 5A, B; also compare with Fig. 2B and D).

The gene *pb* has bilaterally paired strong expression domains (arrowheads, Figs. 4B, 5C), which correspond to the position in which shell adductor muscles are developing (compare Figs. 4B, 5C and 2D). The FISH and CLSM investigation of juveniles further reveals that those two domains extend obliquely from the more anterior point on the dorsal shell to the more posterior point on the ventral shell, in the same orientation as the anterior shell adductors (compare Figs. 5D and 2G).

*Hox3* is expressed in two paired domains posteriorly to the most lateral projections of the shell (arrowheads, Figs. 4C, 5E), where prospective hinge rudiments will form in older juveniles [68]. CISH investigation showed additional broad weak staining in the posterior part of the body (Fig. 4C), which was not reproduced with FISH (Fig. 5E) and which might result from unspecific probe binding in the posterior shell as shown by sense probe staining (Additional file 1, Fig. S1B and C).

The gene *Dfd* is expressed in the ventro-posterior domain (Figs. 4D, 5F–H) composed of extensive lateral elements (arrowheads, Figs. 4D, 5F–H) and a narrow median stripe, detectable only with CISH. Fibrous appearance of those structures in LM imaging (Fig. 4D) as well as their position revealed with FISH (Fig. 5F-H) indicates that *Dfd* is expressed in the ventral pedicle adjustor muscles (compare Figs. 5F–H and 2D, G).

Expression of the gene *Scr* is restricted to the mantle margin (Figs. 4E, 5I). Signal from probes against *Scr* in CISH seems to be diversified into smaller domains with strong signal interspaced by wide regions of relatively weaker expression (respectively: arrowheads and asterisks, Fig. 4E), not visible in FISH examination (compare Figs. 4E and 5I).

The gene *Lox5* is expressed in the continuous dorso-posterior domain, which extend from posterior region of shell-covered body (asterisks, Figs. 4F, 5J) to the pedicle tissues (arrowheads, Figs. 4F, 5J) and its expression is restricted to the dorsal epidermal cells as revealed by FISH (double arrowheads, Fig. 5K).

*Antp* has a distinct expression domain only in the epidermis of the pedicle, as revealed by both CISH (arrowhead, Fig. 4G) and FISH (arrowhead, Fig. 5N; Fig. 5O, P). The signal in the CISH staining developed for the long time (33h in 4°C plus 16h in room temperature) and before it became evident the strong staining had appeared in some specimens also in the dorso-posterior part of the shell-covered body. However, the control with sense probe showed that this staining results from unspecific binding of the probe in larval dorsal protegulum (asterisks, Additional file 1, Figure S1B) and on the borders between dorsal protegulum and remaining parts of the shell (arrowheads, Additional file 1: Figure S1B). The strong dorsal band was also visible in FISH staining (double arrowheads, Fig. 5N) but combined staining with DAPI showed, that it is restricted to the surface area and does not penetrate the epidermis (arrowhead, Fig. S1D, E), supporting our finding that it represents an unspecific probe binding by shell components.

The central-class *Hox* gene *Lox4* is expressed only in the small area of epidermal tissues in the dorso-posterior part of the shell-covered body (arrowheads, Figs. 4H; 5L, M) and its expression domain is not extending to the pedicle tissues.

We did not manage to detect expression of *Post2* with *in situ* hybridization, which corresponds to the reported overall low level of *Post2* transcription in postmetamorphic juveniles of *T. transversa* [19]. After long developmental time (33h in 4°C plus 20h at room temperature) CISH stainings yielded signal in the dorso-posterior part of the shell covered body (asterisk, Fig. 4I), however, the control with the sense probe showed that this results from unspecific binding of the probe in the larval dorsal protegulum (Additional file 1: Figure S1C). The FISH stainings only revealed a signal at the borders of the larval protegulum and remaining parts of the shell (arrows, Fig. S1F) and, similarly as in case of *Antp*, FISH combined with DAPI staining revealed that this signal is restricted to the surface (shell components) and does not penetrate to the cellular epidermal layer (arrowhead, Additional file 1: Figure S1G). The unspecific binding of some probes by the larval protegulum has been already reported for *T. transversa* larvae [79] and apparently this phenomenon can also pose a problem in investigation of postmetamorphic animals.

Expression of the most posterior *Hox* gene *Post1* is detected along mantle margin (arrowheads, Figs. 4J, 5R), showing a relatively equal strength of signal with both CISH and FISH.

## Discussion

### Metamorphosis and *Hox* gene expression in Rhynchonelliformea

Comparison of the expression of *Hox* genes between late, competent larva and postmetamorphic juvenile of *Terebratalia transversa* (Fig. 6) shows that in both stages almost all *Hox* genes (with the exception of *Hox3*, *Post2* and *Post1*) are expressed in corresponding organs and body regions: *lab* in chaetal sacs, *pb* and *Dfd* in, respectively, anterior and posterior mesoderm, *Scr* in the shell growth zone, whereas *Lox5*, *Antp* and *Lox4* are expressed in the dorso-posterior ectoderm. Most of the observed differences and shifts in the expression domains can be explained by the inversion of the mantle lobe, which constitutes the most profound process during the whole metamorphosis in Rhynchonelliformea (Fig. 1B, [63]). Another factor, which contributes to the observed changes, is the restriction of the expression of some *Hox* genes from broad, less specific larval domains to particular structures of the juvenile, which emerge during or after metamorphosis. For example, pb is generally expressed in the anterior mesoderm in late larvae but in juveniles its expression becomes restricted only to particular mesodermal structures, i.e. newly formed anterior shell adductors muscles.

The comparison of the *Hox* gene expression between larvae and juveniles allows to identify Anlagen of adult structures in the larva. For example, the expression patterns of the *Hox* genes before and after metamorphosis suggest that only the posterior half of the larval pedicle lobe contributes to the pedicle of the adult, whereas the more anterior part becomes the posterior region of the shell-covered body, as it has been proposed by Stricker and Reed [63, 71]. Among six *Hox* genes expressed in the pedicle lobe of the late larvae of T. transversa, only *Lox5* and *Antp* are expressed in the pedicle of the postmetamorphic juvenile (Fig. 6), both of them being expressed in the most posterior part of the larval pedicle lobe [19].

Next to Rhynchonelliformea, two inarticulate clades belong to the Brachiopoda: Craniiformea and Linguliformea [80], both possessing a planktonic larvae, which undergoes more or less pronounced metamorphosis [62, 69, 81–85]. In Linguliformea the metamorphosis itself is extended over time with some of the juvenile traits present already in planktotrophic larvae [66, 69, 81, 86] and the most advanced larval stages are even commonly considered as representing planktonic juveniles or paralarvae [62, 69, 86]. One can therefore speculate, that as larval and adult body plans in Linguliformea are continuous, their patterning by *Hox* genes should be similar as is a case in *T. transversa*. On the other hand, there are two competing hypotheses about nature of the rearrangement of the larval body plan during metamorphosis of craniiformean brachiopods [62, 82, 83, 87, 88]. The main controversy regards whether *N. anomala* larva, which lacks the distinct pedicle lobe, attaches to the substrate with its dorso-posterior side [82] or with the posterior tip of the posterior lobe [83]. Expression of the *Hox* genes is relatively similar between embryos and larvae of *N. anomala* and corresponding stages of *T. transversa* [19], indicating a conserved nature of *Hox* genes patterning between Craniiformea and Rhynchonelliformea. *Lox5* and *Antp*, which after metamorphosis are expressed in the pedicle of *T. transversa* juveniles are expressed in the posterior tip of the posterior lobe of *N. anomala* larvae [19] favoring interpretation that posterior tip of *N. anomala* larvae corresponds to the pedicle of Rhynchonelliformea [83, 88]. Further investigation of the postmetamorphic expression of *Hox* genes, especially *Lox4* and *Antp*, in *N. anomala* could support this hypothesis.

Unlike some bilaterians in which metamorphosis seems to be related with highly different *Hox* gene expression between larvae and adults (e.g. tunicates [15], Bryozoa [89]) or in which *Hox* genes are not expressed in the larvae and only pattern adult body (pilidiophoran nemerteans[49], indirectly developing Hemichordates [50], sea urchins [51–53]), rhynchonelliformean brachiopods exhibit continuity in the patterning of larval and adult body plans. Consequently, in regards of *Hox* gene expression, metamorphosis in *T. transversa* is similar to the condition found in another spiralian clade, Annelida. Although there are some shifts in expression patterns of particular *Hox* genes between annelid larvae and juvenile worms [47, 48], those differences are mostly related with restriction of some of the genes from broader larval to more specific adult domains [47]. This similarity can be explained if one assumes that, same as in Annelida, the metamorphosis of rhynchonelliformean larvae is not as drastic as it might seem and instead represents a relatively gradual process [66]. In *T. transversa* several of the adult structures, including shell secreting epithelium [63, 67] or pedicle muscles [69, 71], are already present in the competent larvae as the Anlagen. Thus, even though transition from larva to juvenile poses large ecological change, from the morphological point of view the mantle lobe inversion is related mostly with tissue relocation and not with the degeneration or formation of entire body regions, as is the case in pilidiophoran nemerteans, indirectly developing hemichordates, sea urchins or ascidians.

From the phylogenetic and developmental point of view, it would be interesting to compare shifts of *Hox* genes expression observed during metamorphosis between *T. transversa* and Phoronida. Phoronids are closely related to brachiopods [56, 57, 90, 91] (in past even proposed as specialized clade belonging to Brachiopoda [61, 80]) and their rapid metamorphosis involves drastic rearrangements of the larval body plan [66, 92–95], which is much more complicated than the transition found in Rhynchonelliformea and sometimes referred to as catastrophic or cataclysmic metamorphosis [92, 94, 96]. The recent analysis of the body region specific transcriptomes revealed that in adults of *Phoronis austarlis*, which possesses an organized *Hox* cluster, *Hox* gene expression does not exhibit spatial collinearity [18]. Unfortunately, data on the spatial expression of *Hox* genes in early developmental stages of any phoronid species are still lacking [96], preventing analysis of metamorphosis-related *Hox* genes expression shifts. Nevertheless, it is possible that in phoronids the larvae and juveniles exhibits pronounced differences in the *Hox* genes expression as is a case in some other animals with catastrophic and extensive metamorphosis [15, 49, 50].

### Expression of *Hox* genes during the morphogenesis of brachiopod specific structures in *T. transversa*

Although *Hox* genes are believed to originally be responsible for antero-posterior patterning [1, 2, 11], in certain animal lineages some of them were coopted for morphogenesis of evolutionary novel structures [97–100]. Among Spiralia such phenomenon has been reported in e.g. conchiferan molluscs [36–38] and annelids [34, 35, 47, 48], whereas recently Schiemann et al. suggested that in brachiopod larvae 4 out of 10 *Hox* genes have been recruited for patterning of chaetae (*lab* and *Post1*) and shell fields (*Scr* and *Antp*) [19]. Our results generally support findings of Schiemann et al. – although we did not find evidence for the expression of *Antp* in the shell field – and show that cooption of *Hox* genes for morphogenesis of novel structures is even more pronounced in juveniles of *T. transversa* than it is in the larvae.

*lab* and *Post1* are recruited for the morphogenesis of chaetae in the embryos and larvae of *T. transversa* [19]. The lab is constantly expressed in the chaetal sacs from formation of their early Anlagen up to the latest larval stage, whereas *Post1* is only briefly expressed during short time window, when the Anlagen are formed. In our study we detect expression of *lab* in the chaetal sacs of juveniles as well, but surprisingly we found that *Post1* is also expressed in postmetamorphic juveniles. Moreover, its expression is not only restricted to the chaetal sacs but instead could be detected in the whole marginal zone of the mantle. This finding, however, makes sense when one takes into consideration that as adults T. transversa, as most of the rhynchonelliformean brachiopods, possess numerous chaetae along the whole mantle margin [86, 101]. We therefore propose, that although both *lab* and *Post1* are involved in the chaetae formation in Rhynchonelliformea, they play different roles: *Post1* is expressed in the regions where prospective chaetae will develop, possibly stimulating epidermal cells to differentiate into chaetal sacs before its expression decays. A similar role has been suggested for *Post1* in annelids, whose chaetae are considered homologous to brachiopod ones based on morphological [102] and molecular [19] similarities. In annelids *Post1* is expressed in the cells of developing chaetae-bearing parapodia but the expression faints over the time of development and is not detectable in the already formed parapodia [34, 35, 47]. *lab*, on the other hand, is possibly involved in the patterning of the growth of the chaetae itself, remaining expressed long after onset of chaetal sac formation. This hypothesis needs to be tested in future by functional gene inference and the examination of older juveniles or adults, in which, accordingly, we would expect lack of *Post1* expression and broad expression of *lab* along the whole mantle margin.

The two-valved shell and posterior pedicle represent two distinct apomorphies of brachiopods and we found four out of ten *Hox* genes expressed in the structures related with those morphological novelties. Our results indicates that Scr is likely co-opted for the juvenile shell formation, as the gene is expressed in the mantle margin in the region specialized for shell secretion [68]. This finding corresponds to the results of Schiemann et al. [19], who found expression of Scr in the epithelial cells forming larval shell rudiment. Both shell and pedicle require sets of specialized muscles, which constitute an important part of the brachiopod body. In late larvae of *T. transversa* the genes *Pb* and *Dfd* are likely responsible for A-P patterning of mesoderm [19], yet during post-embryonic development they seem to be recruited into morphogenesis of specific muscular structures that drive the biomechanics of respectively shell and pedicle. Additionally, *Hox3*, another gene that seems to play a role in mesoderm patterning during earlier developmental stages [19], is expressed in the regions where future hinge rudiments will develop [68], suggesting that it could be involved in the morphogenesis of this autapomorphic rhynchonelliformean feature.

## Conclusions

All developmental stages of *Terebratalia transversa*, including juveniles, express *Hox* genes in a spatially non-collinear manner [19]. Most of the patterns observed in the late larvae seem to persist throughout metamorphosis and are retained in juveniles, corroborating morphological observations that metamorphosis, despite being rapid, is of gradual type and most of the adult organs are present as Anlagen in the competent larvae. The most drastic shifts in *Hox* gene expression patterns observed during metamorphosis can be explained by: 1) the inversion of the mantle lobe which relocates some of the more posterior larval structures into the anterior edge of the juveniles and 2) restriction of broad expression domains, present in larvae, to specific structures in juveniles.

Concordantly to the previous study on larvae of *T. transversa* we found that certain *Hox* genes have been evolutionary coopted for morphogenesis of specialized structures in brachiopods. In both larvae and juveniles lab is expressed in the chaetal sacs whereas *Post1* marks the area where prospective chaetae will develop. In juveniles four out of the ten *Hox* genes are expressed in the epidermal (*Scr*, *Hox3*) and muscular (*pb, Dfd*) tissues related with shell and pedicle, two autapomorphic features of Brachiopoda.

## Methods

### Animal collection and fixation

Gravid adults of *Terebratalia transversa* (Sowerby, 1846) were collected near San Juan Island, Washington, USA. Eggs obtained from the animals were fertilized and developing larvae were cultured following previously published protocols (e.g. [19, 63, 67]) up to the metamorphosis. Two days after metamorphosis, juvenile animals were gently scraped from the bottom of the dish with a razor blade, relaxed with MgCl_2_, fixed in 3.7% formaldehyde and washed in phosphate buffer. Fixed animals were stored in 100% methanol.

### In situ hybridization

Probes against *Hox* genes were synthesized using the same plasmid clones as used in Schiemann et al. 2017, where the gene orthology assessment has been performed[19]. Single whole-mount *in situ* hybridization was performed following an established protocol[103]. dUTP-digoxigenin-labelled probes were hybridized at a concentration of 1 ng/μl at 67°C for 48 hours, detected with anti-digoxigenin-AP antibody in 1:5000 concentration in blocking buffer and visualized with nitroblue tetrazolium chloride and 5-bromo-4-chloro-3-indolyl phosphate (in colorimetric *in situ* hybridization) or detected with anti-digoxigenin-POD antibody in 1:200 concentration in blocking buffer and visualized with TSA-Cy5-Plus (in fluorescent in situ hybridization). Additionally, animals prepared for FISH where stained for 30 minutes in DAPI to visualize cell nuclei. Stained juveniles where mounted in 70% glycerol and examined with Zeiss Axiocam HRc connected to a Zeiss Axioscope Ax10 using bright-field Nomarski optics (CISH) or scanned in Leica SP5 confocal laser-scanning microscope (FISH).

### Immunohistochemistry

For investigation of juvenile morphology, mouse primary antibodies against anti-tyrosinated tubulin (Sigma, T9028) and anti-acetylated tubulin (Sigma, T6793) where used in 1:500 concentration. To visualize the primary antibodies, secondary goat anti-mouse antibodies (Life Technologies) conjugated with fluorochrome (AlexaFluor647) were applied in 1:50 concentration. F-actin was visualized with AlexaFluor555-labeled phalloidin and cell nuclei were stained with DAPI. Stained juveniles where mounted in 80% glycerol and scanned in Leica SP5 confocal laser-scanning microscope.

### Image processing and figure preparation

Z- stacks of confocal scans were projected into 2D images and 3D reconstructions in IMARIS 9.1.2. Both light micrographs and CLSM images where adjusted in Adobe Photoshop CS6 and assembled in Adobe Illustrator CS6. All the schematic drawings were done with Adobe Illustrator CS6.

## Declaration

### Ethics approval and consent to participate

Studies of brachiopods do not require ethics approval or consent to participate.

### Consent for publication

Not applicable.

### Availability of data and material

All data generated or analyzed during this study are included in this published article.

### Authors’ contributions

AH designed the study, collected samples and contributed to writing; LG conducted the in situ hybridizations and the confocal studies, arranged figures and drafted the manuscript.

## Acknowledgements

The staff of UW Friday Harbor Laboratories and crew of the vessel “Centennial” are gratefully acknowledged for help in collection of adult *Terebratalia transversa*. We would like to thank Daniel Thiel, Carmen Andrikou and Chema Martin-Duran for help with culturing *T. transversa* larvae and collection of juvenile specimens as well as valuable discussions regarding the project. Additionally, LG is very grateful to Aina Børve for her help and assistance during in situ laboratory procedure.

---

**Fig. S1.**
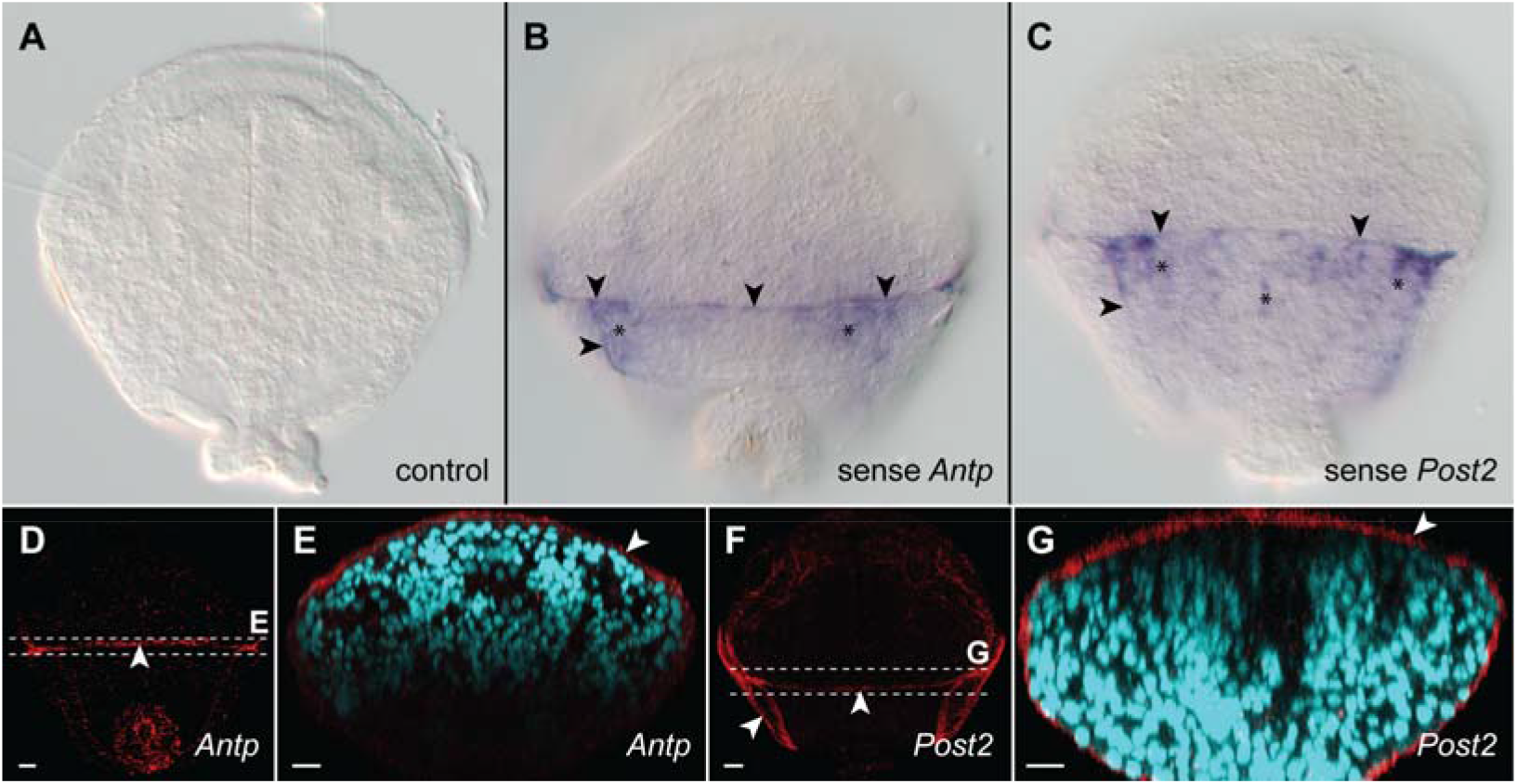
Background signal resulting from unspecific binding of probes by surface of larval dorsal protegulum (asterisks) and borders between protegulum and remaining shell (arrowheads). The control without probes (A). Colorimetric *in situ* hybridization with sense probe of *Antp* (B) and *Post2* (C) genes, signal developed for the same time as for antisense probes. Fluorescent *in situ* hybridization with antisense probes of *Antp* (D, E) and *Post2* (F, G) genes, on E and G combined with DAPI staining of cell nuclei. Dorso-ventral view with anterior to the top (A–D, F) and virtual cross section with dorsal to the top (E, G). Dashed lines with letters on D and F indicate section planes shown on respective plates.

